# Modeling the interplay between plastic tradeoffs and evolution in changing environments

**DOI:** 10.1101/711531

**Authors:** Mikhail Tikhonov, Shamit Kachru, Daniel S. Fisher

**Affiliations:** Department of Physics, Washington University in St Louis, St Louis, MO; Department of Physics, BioX, Stanford University, Stanford, CA; Department of Applied Physics, BioX, Stanford University, Stanford, CA

**Keywords:** Evolution, Changing environments, Performance tradeoff

## Abstract

Performance tradeoffs are ubiquitous in both ecological and evolutionary modeling, yet are usually postulated and built into fitness and ecological landscapes. But tradeoffs depend on genetic background and evolutionary history, and can themselves evolve. We present a simple model capable of capturing the key feedback loop: evolutionary history shapes tradeoff strength, which, in turn, shapes evolutionary future. One consequence of this feedback is that genomes with identical fitness can have different evolutionary properties, shaped by prior environmental exposure. Another is that, generically, the best adaptations to one environment may evolve in another. Our minimal model highlights the need for analysis of simple models capable of incorporating explicit dependence on environment, and can serve as a rich playground for investigating evolution in multiple or changing environments.

---

Performance tradeoffs, caricatured by “you can’t be good at everything”, are ubiquitous in both ecology and evolution. Sometimes modeled explicitly (e.g. as a fixed total budget of energy or proteome), and often implicitly (e.g. as fitness costs associated with traits), tradeoffs are a staple of ecological and evolutionary modeling [1–6].

The rigid tradeoffs assumed in many models implicitly derive from the assumption that long-acting evolutionary pressures drive organisms to approximate Pareto optimality [7–9], i.e. to a regime where performance at the relevant tasks cannot all be improved simultaneously. However, even under this assumption [10], the Pareto front should be high-dimensional [11, 12] so that the tradeoff between any subset of traits is not, in fact, rigid. Furthermore, tradeoffs will depend on evolutionary history and themselves evolve [13–18]. In laboratory experiments, adapting bacteria to one task can both hinder and improve their performance at another, depending on the experimental protocol, the studied strain, and the exact nature of the tasks, as well as history of prior exposure [19–21]. Some phenomena appear non-intuitive and surprising, for instance, even very weak levels of an antibiotic can induce resistance to much higher levels [22, 23].

In some cases, the specific mechanisms responsible for tradeoffs and their plasticity are known, and can be modeled mechanistically [24, 25]. This approach can be very informative about a particular case of interest, but leaves aside important broader questions. Which of the observed behaviors depend on the details of a given biological system, and which are more general? Which experimental observations should be considered surprising, and which can be captured already by the simplest models? What other behaviors might be expected? For instance, can less frequent exposure to environment *X* result in better adaptation to it? Can exposure to *X* result in better fitness in environment *Y* than exposure to *Y* itself? More specifically, would a prior exposure to a milder version of a stress facilitate adaptation to its stronger form?

Elucidating which behaviors are surprising vs. general is a key role of theory and simple models. Here, we propose a minimally structured model capturing some key experimentally observed behaviors: namely, the model exhibits performance tradeoffs, but their strength evolves and depends on evolutionary history. This minimal setting proves sufficient to observe non-trivial ways in which tradeoff strength shaped by evolutionary past can predictably influence evolutionary future; in particular, we identify a mechanism that makes the path towards the highest fitness in one set of environments is via exposure to a different set. Our basic framework can serve as a rich null model for evolution in multiple or changing environments.

## TOOLBOX MODEL

The structure of our model is summarized in Fig. 1A. The environment is represented by a target vector 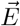 in an abstract *L*-dimensional space, and a genome *G* by a *K*-element basis in that space, with *K* < *L*, i.e. the basis is under-complete. The fitness of genome *G* in environment 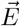 measures how well the basis (the “tools” available) can approximate the target.

**FIG. 1.**
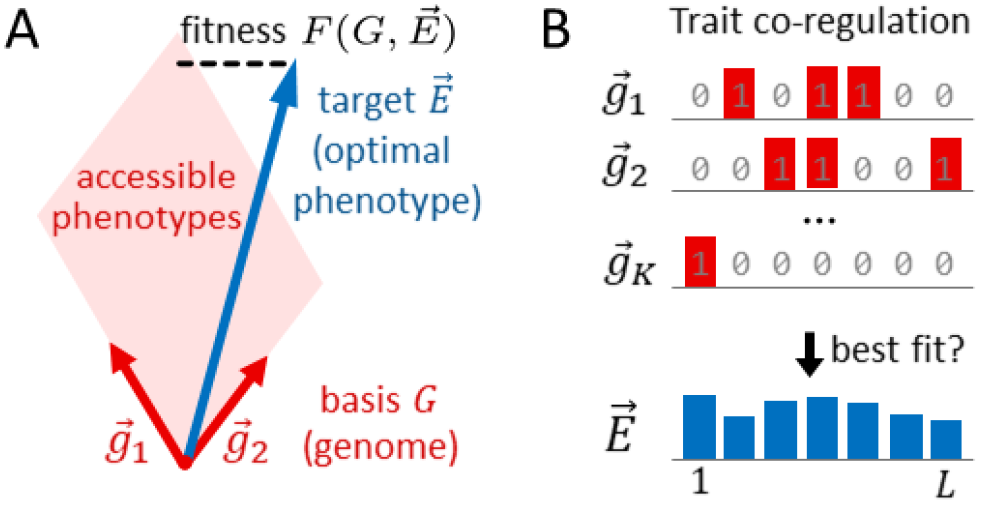
The toolbox model. (*A*) A genome, *G*, consists of an under-complete set of *K* basis vectors 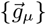 (“tools”). The environment is represented by a target vector 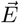. The fitness in this environment is a measure of how well this basis can approximate the target vector, i.e. “fit” the environment. Specifically, 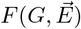 is defined as the residual of the best linear approximation of 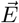 using the vectors 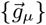 and positive coefficients (“expression levels”). (*B*) For simplicity, we consider 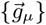 to be binary vectors of length *L*, and assume the components of 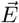 to be positive.

To motivate this setup, consider the *L*-dimensional space as the space of traits (phenotype space). Specifying an environment in our model is equivalent to specifying the optimal phenotype 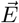 for that environment (here and below, the vector notation always refers to the trait space). An idealized organism with a very flexible physiology, capable of independently adjusting each trait as necessary, would be able to adopt any phenotype. However, an organism limited to only *K* ≪ *L* adjustable “knobs” would in general only be able to approximate the target phenotype. Motivated by this, we caricature a genome by its pattern of trait coregulation, specifically a set of *K* basis vectors 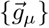, and posit that an organism can adopt any phenotype realizable as a linear combination of its 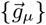 with *positive* coefficients — loosely “expression levels”. Target phenotypes outside this *K*-dimensional subspace can only be approximated, and we define fitness 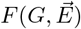 as the (Euclidean) norm of the residual:

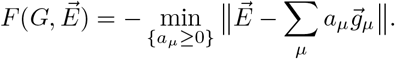

The simplest interpretation of this model is as a caricature of metabolism, where the *L* “traits” are the required amounts of a set of metabolites, and the *K* “knobs” (dimensions of internal representation [26]) are the activities of synthesis pathways. Motivated by this example, we take all components of 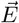 and 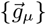 to be positive. However, we stress that our setup is not intended to be a realistic model of metabolic regulation, or any other specific context. Instead, our goal is to construct a minimally structured model that allows the same genome to be good (or not) in many different environments at the same time. In our setup, for any set of *N*_env_ environments with *N*_env_ ≤ *K* there exists, in principle, a genome that is perfectly fit in all. Conversely, the same environment can be fit by many genomes. For our purposes, we will take *N*_env_ ≲ *K* ≪ *L* so that being fit in *N*_env_ environments is possible, but difficult.

Mathematically, the key feature of our model is that the expression levels *a*_*μ*_ are adjusted to be as good as possible in each environment separately. This feature is crucial: An alternative setup, where the values *a*_*μ*_ would be included in the definition of a genome and hence be the same in all environments, is simply a variant of the extensively studied Fisher’s geometric model [27]. Allowing the expression coefficients to depend on the environment – phenotypic plasticity [28–30] – is the key feature that allows the same genetically encoded “toolbox” to be useful in multiple environments.

## MODELING THE EVOLUTIONARY PROCESS

To study tradeoffs, we must consider more than one environment; we focus here on the simplest case: *N*_env_ = 2. Thus the central object is an *environment pair* 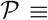 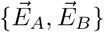. For simplicity we consider binary genomes, where all *L* components of the *K* vectors 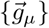 are either 0 or 1 (Fig. 1B); with mutations implemented as bit flips 0 ↦ 1 or 1 ↦ 0. From each genome there are thus *KL* possible mutations.

For simplicity, we analyze the regime of rare mutations and strong selection. In this regime, evolution proceeds through a sequence of sweeps (with no clonal interference), and only beneficial mutations are relevant. As a result, we only need to track the mutations accumulated by a single adapting lineage, avoiding the need to explicitly simulate a population, and eliminating the dependence on population size.

Specifically, given a starting genome *G*_0_ and an environment pair 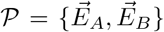, the evolutionary protocol proceeds as follows. One of two environments is randomly chosen. The fitness in this environment of all sigle mutants is evaluated (re-optimizing the expression coefficients every time), and all beneficial mutations identified. Of these, one “lucky” mutation is drawn with probability proportional to its fitness effect (i.e. to its fixation probability). We refer to this as “one mutational step”. After a mutation is accepted, either 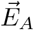 or 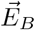 is randomly selected for the next exposure, and the process is repeated. This protocol accepts one mutation per exposure epoch; the validity of this approximation is discussed in the SI (Fig. S1).

## THE TOOLBOX MODEL EXHIBITS TRADEOFFS

We begin by showing that the toolbox model exhibits performance tradeoffs. To do so, we start with *K* = 3 and *L* = 6, small enough that all 2^*KL*^ genomes can be fully enumerated. Panel Fig. 2A shows fitness values *F*_*A*_, *F*_*B*_ of all genomes in two particular random environments: vectors of length *L* = 6, generated by independently drawing each component from an exponential distribution with mean 1, so as to ensure all components are positive. We see that the genomes that perform best in one environment are mediocre in the other, consistent with the notion of a tradeoff. However, even at small K and *L* as shown here, the set of all genomes is too vast to be well-sampled by a single (or a few) evolutionary trajectory, reducing the evolutionary relevance of the global Pareto front (dashed line).

**FIG. 2.**
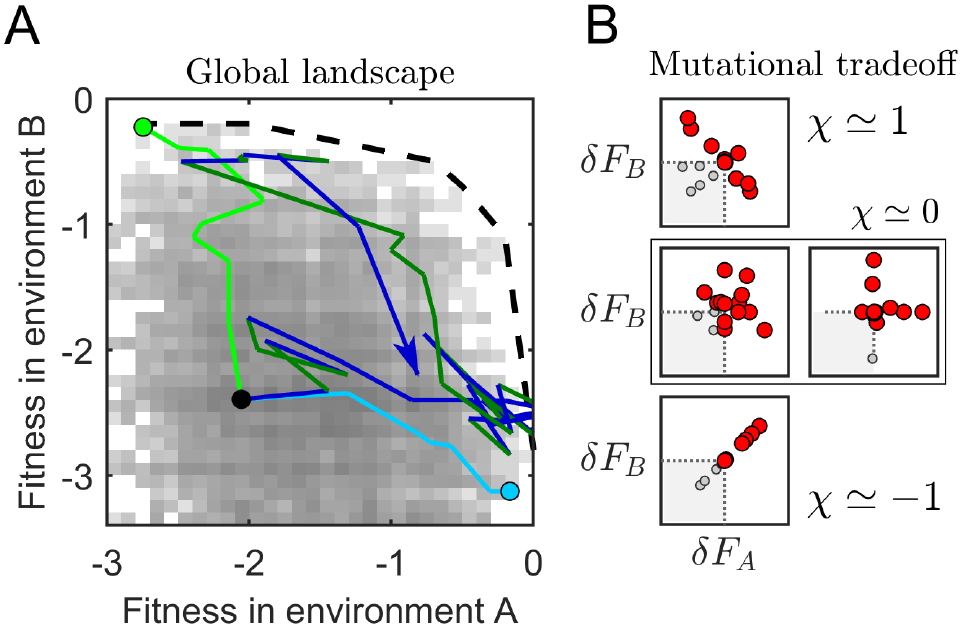
Quantifying tradeoffs. (*A*) The global fitness landscape computed for *K* = 3 and *L* = 6 (small enough that all genomes can be fully enumerated) in two randomly chosen environments 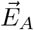 and 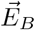. The density scatter plot (gray shading) shows the logarithm of the number of genomes per fitness bin. No genomes are highly fit in both environments and the dashed line roughly traces the Pareto front. A random initial genome evolving under pressure from one environment becomes mediocre in the other (light blue evolving in *A* or light green evolving in *B*) and runs out of beneficial mutations after ~ *L* mutation steps. When the same genome evolves in randomly switching environments, the trajectory does not terminate; steps in dark blue and dark green correspond to mutations accepted while exposed to respectively 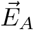 and 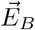 the first 40 steps are shown (some loci flip multiple times). (*B*). Examples illustrating our definition of the *mutational tradeoff χ*. Panels show the relative fitness of all single-step mutants from four example genomes, all far from the Pareto front. In red are mutations beneficial in at least one environment. The doubly-deleterious mutants (gray) cannot fix and are irrelevant for the adaptive evolutionary process studied herein. The examples show: a strong mutational tradeoff (top), the opposite of a tradeoff (bottom); no tradeoff (middle left); and a mutationally modular genome (middle right) where mutations beneficial in one environment have little effect on fitness in the other; see text and Fig. S2.

A more important property for evolution in alternating environments is the *mutational tradeoff* Fig. 2B, on which we will focus. (Other ways of quantifying tradeoffs are discussed in the SI, section “Quantifying tradeoffs”). The scatter plots show the fitness effects of all *KL* = 18 single mutations of several sample genomes, evaluated in the two environments. Mutations deleterious in both environments are irrelevant for the evolutionary process, and can be ignored. The remaining mutations (beneficial in at least one environment) can be used to define mutational tradeoff strength *χ*:

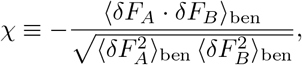

where the subscript indicates the omission of doubly-deleterious mutants in the averaging. This definition ensures that *χ* ranges from −1 (no tradeoff in identical environments: *δF*_*A*_ = *δF*_*B*_) to +1, the strongest possible tradeoff for which *δF*_*A*_ = −*δF*_*B*_. A case that will become important shortly is that of a “mutationally modular” genome, defined by the property that mutations improving performance in one environment do not affect the other (for additional discussion, see SI, “Two definitions of modularity”). Conveniently, by our definition of *χ*, modular genomes have mutational tradeoff of zero (Fig. 2B).

## MUTATIONAL TRADEOFF ITSELF EVOLVES

What are our expectations for the behavior of the mutational tradeoff *χ*? First, any notion of tradeoff strength is expected to depend on the difference between environments. We quantify this difference, Δ*E*, as the component-wise root-mean-square difference between the two target vectors 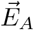 and 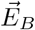. (This definition is more convenient than the Euclidean norm 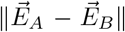 as it ensures that strongly different environments correspond to Δ*E* of order 1 rather than 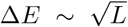.) Second, one might expect the *χ* of an evolving genome to increase with time, since the Pareto front intuition suggests that highly evolved genomes should become depleted for jointly beneficial mutations.

To test these expectations, we investigate the dynamics of mutational tradeoff strength for random initial genomes evolving in random environment pairs with various Δ*E*. As described above, the relevant parameter regime of the toolbox model is *N*_env_ ≲ *K* ≪ *L*. Since our focus is on environment pairs (*N*_env_ = 2), from now on we will use *K* = 4 and *L* = 100. Our starting genomes will be random binary matrices with each entry set to 1 with probability *p* = 0.25 (so that initially, on average, each trait is affected by one regulator: *pK* = 1). For each trial, the components of both environment vectors are independently drawn from an exponential distribution as before (using a Gaussian distribution does not qualitatively change the results), and the component-wise differences are then rescaled to achieve a desired Δ*E* between the pair (see SI, section “Parameterizing environment pairs”). The results are presented in Fig. 3A.

**FIG. 3.**
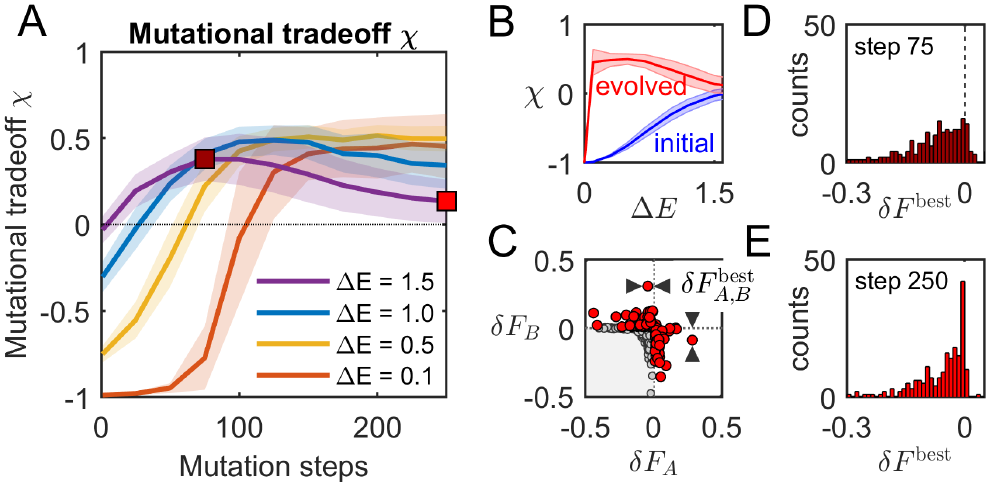
Evolved tradeoffs are not typical. (*A*) Mutational tradeoff (*cf.* Fig. 2B) as a function of the number of mutation steps (a proxy for time), for evolution in random environment pairs differing by a given amount, quantified by Δ*E*. Each curve shows mean ± 1 s.d. over 100 independent trials (shaded); for each trial, a new random initial genome was evolved in a new random environment pair (*K* = 4, *L* = 100). Initially, more different tasks are associated with stronger mutational tradeoff, and tradeoff strength generally increases with time, with a notable exception at large Δ*E*. (*B*) Tradeoffs after 250 mutational steps versus environment-difference Δ*E*. For Δ*E* = 0 (two identical environments), the mutational tradeoff is necessarily −1. However, when the environments differ, evolution drives tradeoff strength to strongly atypical values. Shown is mean ± 1 s.d. over 100 independent trials for each Δ*E*. (*C*) The mutant cloud for an example genome evolved for 250 steps in two very different environments (Δ*E* = 1.5). Apparent modularity emerges: the best mutant in one environment has only a weak effect (denoted 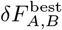) in the other. (*D, E* ) Histograms of *δF*^best^ over 100 trajectories evolved at Δ*E* = 1.5, at two timepoints highlighted in *A*. The later timepoint *E* shows a clear enrichment of modular genomes (peak at *δF*^best^ ≈ 0).

Plotting these data as a function of Δ*E* (Fig. 3B) appears to confirm much of our intuition. First, for random genomes (timepoint 0), mutational tradeoff *χ* is strongest when Δ*E* is largest: as expected, being good at two tasks is harder when they differ more. Second, genomes evolved for 250 mutation steps exhibit the expected increase of the mutational tradeoff. For a sense of scale, recall that our genomes have *KL* = 400 loci, so 250 steps are sufficient for over half of the bits to flip.

Curiously, however, Fig. 3A also shows that after ~ 100 mutation steps, genomes evolving at a large Δ*E* consistently exhibit a counter-intuitive *decline* in *χ*. This behavior is not a peculiarity of our definition of mutational tradeoff; other measures of tradeoff strength exhibit the same phenomenon (Fig. S3). Examining the evolved genomes in more detail reveals that this decline in *χ* is associated with genomes becoming more modular: Fig. 3C shows the mutant cloud for one of the genomes evolved at Δ*E* = 1.5. Its “plus”-like shape is indicative of emergent modularity (cf. Fig. 2B); in particular, the best mutant in one environment has only a weak fitness effect in the other. Denote the cross-effect of the two best mutants (one for each environment) as 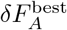 and 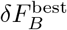. A mutationally modular genome is one with 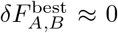. Panels D and E show histograms of *δF*^best^ observed in 100 independent replicates (random starting points, random environment pair with Δ*E* = 1.5). Panel E (the later timepoint) shows a clear enrichment of modular genomes (the peak at *δF*^best^ ≈ 0), compared to panel D (the earlier timepoint).

One explanation for this enrichment might be that high-fitness genomes are generally more modular. However, we will now demonstrate that evolution at a large Δ*E* specifically promotes tradeoff weakening: even conditioned on having the same high fitness, the *χ* of evolved genomes remains atypical [31]. Specifically, we will show that (a) high-fitness genomes can exhibit a whole range of mutational tradeoff values; (b) prior evolutionary history predictably pushes genomes into different regions of this high-fitness space, and (c) the resulting genomes, although sharing the same fitness, can differ dramatically in their properties (for instance, in their ability to evolve further).

## EVOLUTIONARY HISTORY SHAPES MUTATIONAL TRADEOFF

Fig. 3B demonstrated that mutational tradeoff strength of evolved genomes is not typical of all genomes (the red curve is clearly distinct from the blue). Establishing whether evolution specifically promotes mutational tradeoff weakening requires a stronger statement, namely that the *χ* arising through evolution is not even typical of high-fitness genomes. To show this, the most direct approach would be to compare the evolved *χ* to the typical values observed when sampling high-fitness genomes in an unbiased way. Unfortunately, we have no procedure for such an unbiased sampling, other than a complete enumeration which is only viable for extremely small *K* and *L* (cf. Fig. 2). Instead, we will reach the same conclusion by showing that different ways of evolving high fitness lead to genomes with different mutational tradeoff strength.

Consider the following computational experiment. Generate one random environment pair 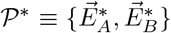 with Δ*E* = 1 for concreteness. Define *fitness in the environment pair* as the average over fitness in 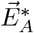 and 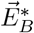:

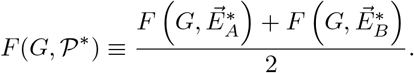

Throughout the experiment, we will only be concerned with fitness and mutational tradeoff as measured in this pair 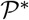. Under our protocol, the fixation probability of any mutation depends only on its fitness effect in the one environment to which the genome is exposed at the time. However, as exposures alternate, the average fitness in the pair will typically also increase.

What other evolutionary protocols could lead to increased fitness in 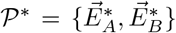? One cannot expect that evolving in some random other pair will increase the mean fitness in 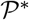, but one can consider evolving a genome in similar environments, or the average of 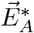 and 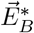. In our model, for any 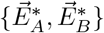, one can create similar pairs of environments that are closer or further apart by linear interpolation or extrapolation (in log space to preserve positivity of components; see SI). This yields a one-parameter family of environment pairs 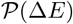 indexed by their difference Δ*E* (Fig. 4A). For a concrete analogy, if 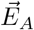 and 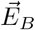 represent hot and cold seasons, then 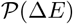 is a family of environment pairs where the intensity of seasonal variation is more mild or more severe (quantified by Δ*E*). If we pre-evolve the same 10 random starting genomes in these conditions, with Δ*E* exaggerated or softened as described, what effect will this have on the fitness and mutational tradeoff strength as measured in the pair of interest, 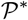?

**FIG. 4.**
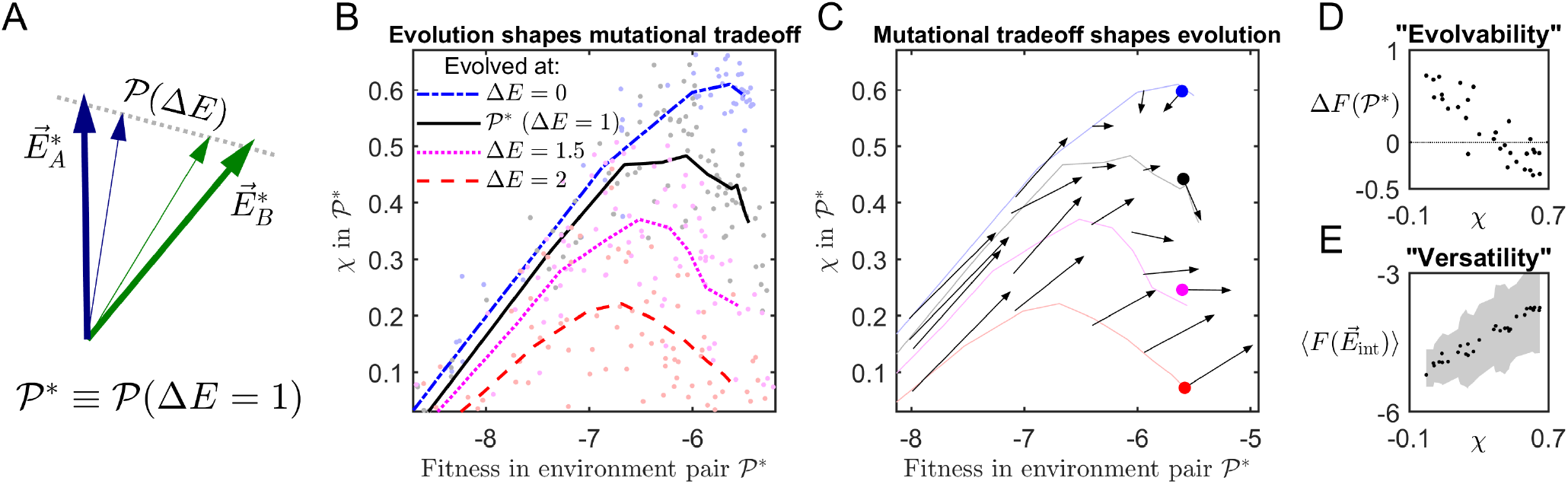
Evolutionary history shapes tradeoff structure, and vice versa. (*A*) For a randomly chosen pair of environments 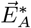, 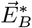, linear interpolation or extrapolation (in the log space, see SI) is used to exaggerate or soften the difference between them. This family of pairs is denoted 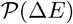, with 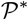 the original pair which differ by Δ*E* = 1. (*B*) The same 10 random initial genomes (*K* = 4, *L* = 100) were evolved either directly in 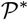(solid black line), or in another environment pair from the 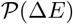 family, with Δ*E* = 0 (the average environment) or Δ*E* = 1.5, 2 (exaggerated differences). The mutational tradeoff measured in the original environment pair 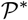 is plotted against the mean fitness in 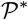: i.e., genomes are evolved in different environment pairs, but are all evaluated in the same pair. Data points show the individual genomes every 25 mutational steps; each line is the average over the 10 trajectories evolved in the same environment pair. We observe that different evolutionary histories consistently drive genomes to different tradeoff strengths, even when compared at the same fitness. (*C*) Mutational tradeoff shapes the near-term evolutionary future. Here, sets of 10 genomes, prepared by pre-evolving for varying lengths of time at different Δ*E* as in panel B, were “transferred” to continue evolution in the pair 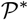. For each set, an arrow shows the average change of fitness and mutational tradeoff in 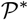 over the next 20 mutation steps. Trajectories from panel B are shown for reference. The contrast in outcomes is particularly striking for the highest-fitness genomes (colored dots); see also next panel.(*D*) “Evolvability” of genomes with different mutational tradeoff. From each of the 4 × 10 trajectories described in panel B, we selected the genome whose mean fitness in 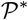 was closest to −5.6 (within ±0.01), yielding 31 genomes (9 trajectories did not attain this high a mean fitness). Each was then evolved in 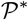for 20 steps; the mean-fitness gain (over the two environments in the pair 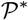) is plotted against the initial value of mutational tradeoff *χ*. The genomes with the weakest *χ* evolve fastest, whereas genomes with the highest *χ* actually decline in mean fitness (compare with C). (*E*) The most evolvable genomes are also least “versatile”. Same 31 genomes as in D, evaluated in random “intermediate” environments 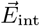 between 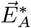 and 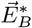; see text. Shown are means ± 1 s.d. over 100 intermediate environments.

The results are presented in Fig. 4B; showing the mutational tradeoff versus average fitness, both measured in 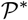. The black solid line corresponds to the simplest scenario, where the 10 initial genomes are evolved directly in 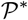. The evolutionary time runs left to right, reflected in increasing fitness. If our hypothesis is correct, evolving the same initial genomes at a larger Δ*E* should promote a more modular genome architecture and thus a weaker tradeoff; this is indeed what we observe (Fig. 4A; magenta and red). Conversely, if instead of exaggerating the differences we soften them to Δ*E* = 0 (i.e. replace the environment pair 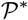 with a single environment, their mean), we obtain the trajectory in blue. (Note that the mutational tradeoff evaluated in an environment “pair” with Δ*E* = 0 is always −1, but panel Fig. 4B shows the mutational tradeoff evaluated in the original pair of interest 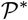)

Fig. 4B directly demonstrates that, within our model, manipulating evolutionary history predictably pushes evolving genomes towards stronger or weaker mutational tradeoff. In particular, if we pick similarly-performing genomes from the right-hand side of this plot, we will find that they all attain the same mean performance in different ways, with a wide range of tradeoff values. These differences must clearly have consequences for their evolutionary properties. Since by the process of evolving in pairs of environments whose differences are exaggerated or softened one can generate genomes anywhere on the mean-fitness vs. mutational tradeoff plane, we can ask: how does the near-term evolutionary future differ for genomes starting at different points of this plane?

## TRADEOFF STRENGTH SHAPES EVOLUTION

To address how the evolutionary history and current tradeoff strength affect future evolution, we used the protocol of Fig. 4B (computationally pre-evolving the same 10 initial genomes in different environment pairs for a varying length of time) to obtain a collection of 10-genome sets evenly populating a region of the fitness vs. mutational tradeoff plane. Each such 10-genome set was then evolved for 20 further mutational steps in our pair of interest 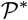. The result is presented in Fig. 4C, where each arrow describes the change in the mean fitness and tradeoff strength in 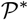 observed over the 20 mutation steps (one arrow per 10-genome set).

First, notice that arrows in the vicinity of the black trajectory are tangent to it. This, of course, is exactly what we expect, since the black trajectory (copied from Fig. 4B) traces the evolution in this same pair 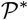. However, arrows starting elsewhere on this plane (pre-evolved at a different Δ*E*, then transferred into 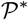) show different behaviors. It is especially interesting to compare the arrows on the far-right side of the plot. And as a reminder, under our protocol, these four 10-genome sets (colored dots) were obtained from the same 10 initial genomes and differ only by evolutionary history. Yet evolving them in the same environment 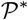 for the same amount of time (20 mutation steps) leads to very different outcomes. The low-tradeoff genomes obtained by pre-evolving at an exaggerated Δ*E* (red dot) exhibit a dramatically faster speed of subsequent evolution in 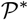, compared to similar-fitness genomes that never left this environment pair (black dot). In contrast, attempting to further evolve the highest-tradeoff genomes (blue dot) actually leads to a *decrease* in mean fitness. For these genomes, mutations beneficial in one environment of the pair are so strongly deleterious in the other that the mean fitness declines. To summarize, genomes with different evolutionary history have predictably different evolutionary future.

This observation is further illustrated in panel D, where the mean-fitness gain over 20 generations is shown for 31 individual genomes (all with the same initial mean fitness −5.60 ± 0.01), plotted against their initial mutational tradeoff strength. The plot directly confirms that weak-tradeoff genomes evolve fast, whereas for the strongest-tradeoff genomes fitness gain dips into the negative.

The relation observed in Fig. 4D is consistent with the previously proposed idea that modular architecture is more “evolvable” [32–34]. Intriguingly, however, our framework allows the benefits of modularity to be nuanced. For example, rather than continuing evolution in the pair 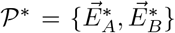, consider instead generating a large number of “intermediate” environments 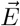, where each component is independently and uniformly drawn to be between the respective component of 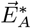 and 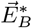. Fig. 4E shows the average fitness of the exact same 31 genomes in the intermediate environments. The trend is now reversed: the “best” genomes as judged by panel D perform worst in this test. Intuitively, while a modular architecture (for a given environment pair) facilitates continuing adaptation to that same pair (Fig. 4D), it also suffers from a form of “overfitting”: the non-modular architectures may be more versatile when evaluated across a range of similar environments (Fig. 4E).

In summary, Fig. 4 demonstrates that evolution shapes mutational tradeoff, and conversely, mutational tradeoff shapes evolution. We will conclude by exhibiting a qualitatively nontrivial phenomenon arising as a consequence of this important feedback loop.

## BEST ADAPTATION FOR ONE ENVIRONMENT PAIR EVOLVES IN ANOTHER

Consider the problem in which a pair 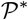 is given, and we would like to evolve a random starting genome *G*_0_ towards a high mean-fitness in this particular pair of environments. The most natural approach, of course, would be to evolve *G*_0_ directly in 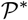. However, the results of Fig. 4 suggest that evolving instead in a different environment pair, with differences exaggerated or softened, might offer a more efficient path towards 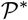-fitness increase. Fig. 5A confirms this by re-plotting the trajectories from Fig. 4A, showing fitness in the environment pair of interest 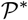 (with Δ*E* = 1) as a function of time, for genomes evolving under 3 different values of Δ*E*. Importantly, in this example, genomes evolving directly in 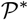 are typically *less* fit than those evolved from the same initial genomes, and for the same number of mutation steps, but in a different environment pair (blue or red).

**FIG. 5.**
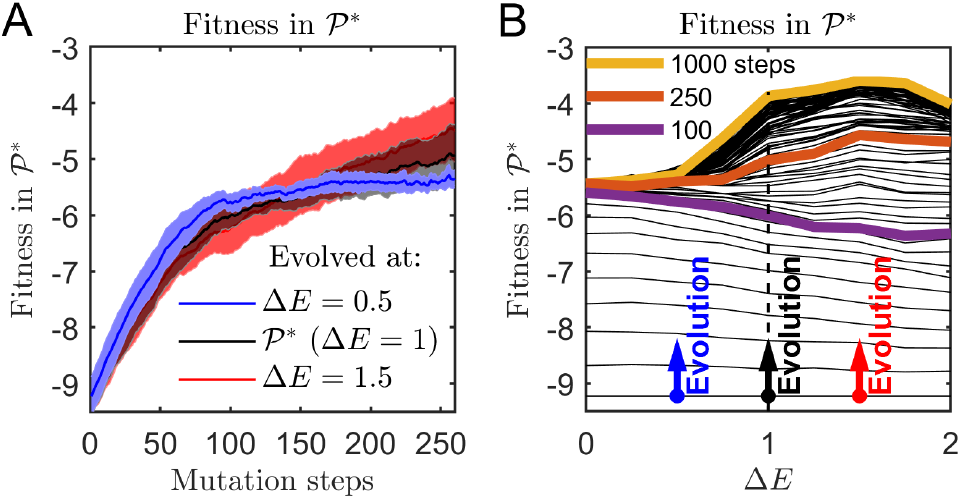
Best adaptation for one environment evolves in another. (*A*) The same 20 random initial genomes (*K* = 4, *L* = 100) are evolved at different Δ*E*, and evaluated at Δ*E* = 1. Shown is mean fitness ± 1 s.d. over the 20 trajectories. Remarkably, the genomes evolving directly in the environment pair of interest (black) are typically less fit than those evolved from the same initial genomes, and for the same number of mutation steps, but in one of the other environment pairs with softened (blue) or exaggerated (red) differences. (*B*) Same as (A), replotted versus Δ*E*. The same initial genome is evolved in different environment pairs; contour lines spaced by 10 mutation steps show the mean fitness evaluated in the environment pair of interest (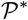 with Δ*E* = 1). Plot averaged over 20 random instances of the initial genome. The three arrows correspond to the three trajectories shown in panel A. We see that generically, the genomes with the highest fitness at Δ*E* = 1 were evolved in *other* environment pairs: after 100 mutation steps, the genomes performing best at Δ*E* = 1 are those evolved at Δ*E* = 0, while at 250 muta-tion steps, the best were evolved at Δ*E* = 1.5. See text for explanation.

A more detailed visualization of this phenomenon is presented in Fig. 5B. In the hot-and-cold season analogy, think of the Δ*E* axis in panel Fig. 5B as a transect through a continent, running from a region with no seasonal variation (Δ*E* = 0) through regions where it is increasingly more severe; we are focussing on a particular temperate region 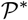 with moderate seasonal variation (Δ*E*\* = 1). Imagine populating this continent with an initially clonal population (the initial genome *G*_0_). Initially neglecting migration, we let these genomes independently evolve in their respective environments. However, every 10 mutation steps, we consider potential immigrants to the temperate region of interest: namely, we evaluate the performance of all the lineages in the pair 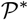 and plot it on the *Y* axis.

By construction, initially all genomes are the same, and their performance in 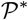 is also the same. However, 100 mutation steps later (purple), the highest fit-ness in 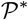 is exhibited by genomes that evolved at Δ*E* = 0. These genomes were evolved under selection pressure from a single, averaged environment and cannot, of course, develop any hot- or cold-specific adaptations, but at the early stages, evolution proceeds very efficiently as there are no conflicting pressures from the two environments.

As evolution proceeds, the behavior inverts. After 250 mutation steps (orange), the highest fitness in 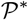 is exhibited by genomes that evolved at Δ*E* ≈ 1.5. Initially, the increase in mean fitness under such harsh seasonal variations was very slow, with beneficial mutations in one environment undoing the gain made in the other. However, eventually the lineages evolving under this large Δ*E* develop a weakened mutational tradeoff (cf. Fig. 4): this enables them to gain fitness more efficiently, overtaking the lineages evolving in 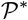. In this model, modularity is thus good for evolvability, but takes time and the right conditions to itself evolve.

It is worth noting that in this situation with a continuum of possible pairs of environments, the lineages evolving directly in the environment of interest are not, in fact, privileged: one generically expects the best fitness to be achieved at a different Δ*E*, even in the long run (shown in yellow for 1000 steps). One implication of this in our spatial / seasonal metaphor is that in the presence of migration, the phenomenon described will lead to a qualitative change in the expected genealogy structure of the long-term surviving lineages. In particular, the environment pairs with large variations turn into sources of evolutionary innovation, promoting the development of 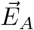- and 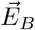-specific adaptations capable of invading other environments and out-competing the resident lineages. Importantly, in our model this is true even though we have explicitly excluded the effects of diversity that would in general be created in each environment.

## DISCUSSION

Understanding evolution in multiple or changing environments requires developing an understanding of which phenomena observed in the laboratory (or the wild) depend on specific details (at molecular, cellular, or higher order level), and which are more general; which experimental outcomes are truly surprising, and which can be found in a very simple model. We presented a toy model able to capture the key feedback loop of evolutionary tradeoff plasticity, whereby organisms evolving in different environments are constrained by performance tradeoffs, but such tradeoffs themselves depend on the evolutionary history. By defining fitness through a regulatory or physiological optimization problem, our approach is reminiscent of asking evolutionary questions from within the framework of flux balance analysis [35–41]. However, our “toolbox” model abstracts away any specifically metabolic (or other) detail, retaining only the flexibility of regulation, i.e. the fact that genome-encoded tools can be used in an environment-dependent manner. Remarkably, this simple model already exhibits qualitatively rich phenomena, including effects that change sign during the course of evolution.

One of the effects we studied by example shows how the best adaptation for one environment may be expected to first emerge in another. Examples of this phenomenon are known experimentally: for instance, the fastest way to evolve resistance to a high dose of antibiotic is through a series of exposures to increasing doses, rather than direct pressure from the environment of interest (see also [19, 23]). Our results suggest that for evolution considered across multiple environments, this scenario may well be generic. If true, this is likely to impose a strong constraint on the predictive power of single-environment evolutionary models: successful lineages coming from elsewhere are beyond their scope, yet successful invaders can have a profound impact, including on genealogical structures, a key observable used for inferences from data. This highlights the need for simple illustrative models able to incorporate explicit dependence on environment. We hope the model presented here will help develop null-model phenomenological expectations, against which experimental (and genealogical) results can be compared (e.g. [21]).

The phenomena reported here did not require finetuning of model parameters; they arise because the space of high-fitness genomes is naturally large and diverse. As a result, different evolutionary histories generically bias evolved genomes towards different corners of the high-fitness space. In our model this property is ensured by choosing the number of tools that can be independently regulated to be larger than the number of environments probed (*K* > *N*_env_), but the observation that there are many ways to be fit should surely not be model-specific. Establishing the generality of our observations, extending them to multiple environments (rather than just pairs), and relating the predictions of our framework to experiments, all constitute productive directions for future work. Furthermore, our simple model assumed organisms sense their environment perfectly and regulate their physiology optimally. Relaxing these assumptions offers natural directions in which our model could be extended.

## ACKNOWLEDGMENTS

We thank A. Agarwala, S. Leibler, and R. Monasson for helpful discussions. This work was supported in part by National Science Foundation grants PHY-1607606, PHY-1545840, PHY-1720397 and a Simons Investigator Award to S.K. All simulations performed in MatLab (Mathworks, Inc). The associated code, and scripts reproducing all the figures in the paper are available at http://doi.org/10.17632/ykypdppy9n.1

## SUPPLEMENTARY INFORMATION

### S1. THE TOOLBOX MODEL

The toolbox model defines a genome as a set of “tools” (basis vectors), whose ability to approximate a given target is interpreted as fitness. The two key parameters here are *K* (the number of basis vectors) and *L* (the length of these vectors). In this section, we discuss the role of these parameters, as well as the one-mutation-per-exposure-epoch assumption of our evolutionary protocol, and the choice to parameterize environment pairs by their difference Δ*E*.

#### *K* sets the largest number of environments where a genome can be fit

In the toolbox model, *K* is the dimensionality of the phenotype space a genome can access. We described it in the main text as the number of regulatory knobs an organism has at its disposal. Thus *K* specifies the largest number of “independent” environments in which a genome can be perfectly fit.

In more formal terms, imagine we draw *M* target vectors, and ask if there exists a genome perfectly fit in all *M*. To the first approximation, the answer is that such a genome would exist if and only if *M* ≤ *K*. (This answer would be exact if we did not require genome entries to be binary, and expression coefficients to be positive.)

Note that the existence of a perfect genome is a different question from whether it would emerge through evolution, or what path the evolutionary trajectory would take. For much of this work, we considered genomes with *K* = 4 evolving in environment pairs. In these circumstances, the question of whether good genomes exist is trivial: in fact, the space of high-fitness genomes is guaranteed to be large. It is precisely this property that makes it possible for evolutionary history to shape the outcome in interesting ways: since the space of high-fitness genomes is large, a genome’s history can influence which corner of this space it will come to occupy. Thus, for the purposes of this work, we want *K* to be larger than the number of environments *N*_env_. At the same time, it should not be *much* larger, as *K* ≫ *N*_env_ would make accommodating *N*_env_ environments too easy (no tradeoffs). This is why, as stated in the main text, the ideal regime for uncovering interesting phenomenology of tradeoff evolution is *K* ≳ *N*_env_. Incidentally, this discussion shows than for a fixed *K*, one may expect a qualitative change in behavior as a function of *N*_env_ in the vicinity of *N*_env_ ≃ *K*, highlighting an interesting avenue for future work.

#### *L* and the frequency of environment switching

The main text introduces *L* as the dimensionality of the trait space. The role played by this parameter can be understood as follows. In a given environment, it will take of order *L* mutations to substantially improve fitness (and if the environment is held static, it will take ~ *L* mutation steps for a genome to run out of beneficial mutations; cf. Fig. 2A). If *L* is small, even a single mutation while exposed to environment 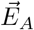 may substantially hurt the performance in the other, 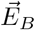. In contrast, when *L* is large, the fitness effect of any one mutation is small.

If we focus on expression coefficients rather than fitness, the situation is similar: After each mutation, the optimal expression coefficients are determined anew, but for a large *L* and a genome close to random the change of optimal expression is adiabatic, with each mutation inducing a change of order ~ 1/*L*. Thus the timescale for any significant change in genome architecture, e.g. a basis vector becoming specialized to environment 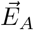 (with a close-to-zero expression when exposed to 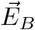), is again *L* mutations. During this time the organism will have experienced *L* environment switches, so a larger *L* corresponds to more switches over the relevant timescale. For instance, the simulations for Fig. 3–5 in the main text used *L* = 100 and all trajectories are shown on the scale of ~ 100 mutation steps, while environment has a 50% switching probability after each. To study the slow-switching regime, one could decrease *L* and/or increase the number of mutations accepted at each exposure.

#### The one-mutation-per-exposure-epoch assumption

As shown above, the limit of *L* → ∞ effectively corresponds to increasingly frequent environment switching. However, we should stress that this switching never becomes “infinitely fast”: by construction, every exposure is long enough for a mutation to occur. We see this as a feature, not a bug. One might have imagined that if environments are switched sufficiently rapidly, one should recover evolution in a *single* environment, namely the average of the two. However, this limit is poorly defined, since such high switching frequency violates the approximation of an environment-defined fitness landscape, and cannot be accessed from within a model using this term. In addition to simplifying the simulations, the one-mutation-per-exposure-epoch assumption made in our model prevents us from accessing this ill-defined regime: even the fastest-switching regime remains just slow enough to allow calling our protocol as “evolution in a pair of environments”. We thank A. Murugan for helpful discussions on this topic.

For all its benefits, this assumption is an approximation, and we should comment on its validity. First, here is an example where such an assumption would be disastrously incorrect. Imagine a genome *G*_0_ that has plenty of room for improvement in environment 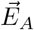, but is almost perfect in environment 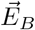; in the former, many strong beneficial mutations are available; in the latter, let’s say there is only a single beneficial mutation *μ*\* with a very small fitness effect. In these circumstances, we should expect many mutations beneficial in 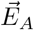 to fix before *μ*\*. Under our protocol, however, the probability of *μ*\* to fix by the next step is 50%.

This example illustrates how the validity of our approximation should be assessed. Denote {*μ*_*A*_} the beneficial mutations available in environment 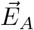, and similarly for environment 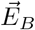. If, forgoing approximations, we were to simulate a sequence of short exposure epochs of length *τ* (short enough that many would elapse without any mutation fixing, long enough that any selection sweep can still be considered instantaneous), what is the probability that the first fixed mutation will be from the set {*μ*_*A*_} (as opposed to {*μ*_*B*_})? In the limit of rare mutations, each mutation, when arises, will escape drift with probability proportional to its fitness effect, at which point it will deterministically sweep through the population and fix. The probability that the next fixed mutation will be from the set *μ_A_* is thus given by

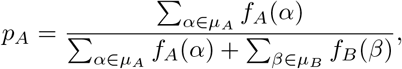

where *f*_*A*_(*x*) and *f*_*B*_(*x*) are the fitness effects of mutation *x* in environments 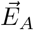 and 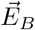, respectively.

Our approximation is to assume that the availability of beneficial mutations in the two environments is balanced, and *p*_*A*_ ≈ 1/2. Since the initial genomes and environments are all drawn randomly, one expects this to be a good approximation at least at the initial stages. Let us examine its validity for the longest time-trajectories in this work, namely the simulations of Fig. 5 (up to 1000 mutation steps).

Instead of simply comparing *p*_*A*_ to 1/2, it is more informative to look directly at the “total mutation potential” in the two environments: 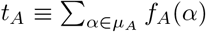 for environment 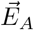, and similarly for 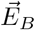. Our approximation is valid as long as *t*_*A*_ ≈ *t*_*B*_.

Fig. S1 plots *t*_*B*_ versus *t*_*A*_ for the three sets of evolutionary trajectories in Fig. 5A. A random genome has a low fitness and strong beneficial mutations are plentiful. (Of course, the effects of mutations are always evaluated in the environments where a genome evolves.) As time goes on, fitness improves and availability of beneficial mutations decreases. (We measure time in mutation steps, but in absolute units each step becomes increasingly longer.) Still, for mutation steps up to ~ 250 (purple, orange) used through most of this work, the two environments remain well balanced, and our one-mutation-per-exposure epoch approximation is justified. Beyond 250 mutation steps (yellow), we are beginning to observe periods of strong imbalance, alerting us that our approximation is starting to fail: if strong mutations in one (and only one) of the environments are depleted, our protocol confers an unfair advantage to the weak ones. We must, therefore, stress that none of the conclusions of this work relied on this long-time regime: these extended simulations were used only in Fig. 5B and exclusively for illustration purposes, to highlight that even in the long term, evolving directly in the environment pair of interest is not guaranteed to be the best strategy to achieve highest fitness. All the phenomena discussed in this work were observed in the early stages of evolutionary trajectories, a regime where our simplifying approximation remains valid.

**FIG. S1.**
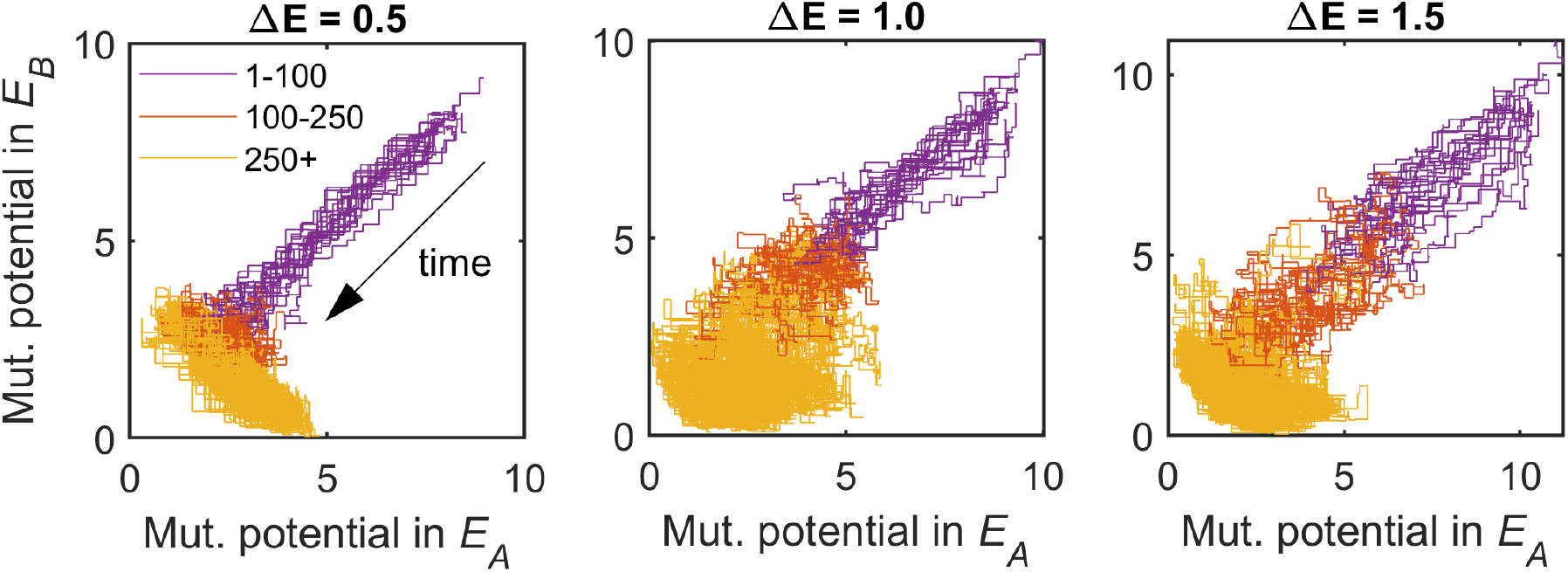
The total mutation potential (sum of effects of all beneficial mutations available) in the two environments in which a genome is evolving, for the trajectories of Fig. 5A in the main text. The left, middle and right panels correspond to respectively the blue, black and red traces on Fig. 5A. Colors as in Fig. 5B: first 100 steps in purple, steps 100-250 in orange, steps 250-1000 in yellow. The plot confirms that through the first ~ 250 mutation steps, the availability of beneficial mutations in the two environment remains balanced (trajectory follows the diagonal). This justifies our procedure of accepting one mutation per environment exposure epoch; see accompanying SI text.

#### The role of initial genome density *p*

Our evolutionary simulations also use a parameter *p*. Unlike *K* and *L*, the parameter *p* describes not the model, but the initial state: it is the probability with which we set each genome entry to 1 at the initial timepoint (the initial genome density). Intuitively, *p* connects regimes where each trait is affected by at most a single regulator (small *p*) and one where “everything affects everything” (large *p*). For all our simulations, for the sake of concreteness we set *p* = 1/*K* = 0.25 (each trait affected by one regulator on average). We did not systematically vary this parameter or explore its effect. As soon as the evolutionary process begins, the genome entries are free to change as they please, so this initial density is not maintained over time. However, it may still have biasing effects on evolutionary future. Intuitively, one might perhaps expect that a low-density starting genome may be more likely to evolve into a modular architecture, while a high-density starting point might favor “generalist” genomes, with all basis vectors used in all environments. We leave these questions for future work.

#### Parameterizing environment pairs

Figs. 4 and 5 in the main text refer to a procedure for “exaggerating” or “softening” the differences between the two environments in a given pair (Fig. 4A). The simplest way to implement it would be to define 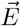 as the arithmetic mean 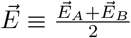, and posit:

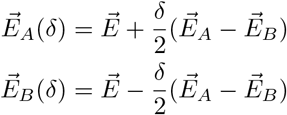

This defines a one-parameter family of environment pairs (indexed by *δ*), where *δ* = 1 returns the original pair, and *δ* = 0 reduces it to a single environment (their mean). The only issue with this approach is that at *δ* > 1 (when differences are exaggerated), some components of the environment vectors may become negative.

In principle, allowing negative components in the target vector (while still requiring expression coefficients to be positive) would merely impose an extra fitness penalty on every genome. Therefore, we could have adopted the procedure above, and the results would be qualitatively similar. However, for consistency, we opted for a protocol preserving the positivity constraint, applying the same procedure as above, but in log space:

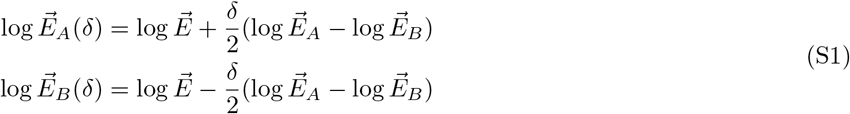

where logarithms are applied to each vector component, and 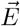 is now the *geometric* mean (=arithmetic mean in log space). (S1) is the simplest procedure that respects the positivity constraint, and was used throughout this work. The indexing by *δ* is trivially converted into indexing by Δ*E*, since Δ*E*(*δ*) is monotonically increasing and thus invertible. The procedure we just described seems quite natural, but it is by no means unique or privileged (as already apparent from the discussion above, since the choice to use linear or logarithmic scale was rather arbitrary). Our goal was to demonstrate that different ways of obtaining highly fit genomes result in genomes differing in tradeoff strength and other evolutionary properties; any other family of evolutionary protocols supporting this observation would have been an equally good choice for Fig. 4 and 5.

Separately, an interesting more general question worth considering is: what features of the 2*L*-dimensional space of environment pairs 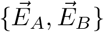 matter for an evolving genome? In the limit of large *K* and *L*, the problem of positive-coefficient fitting of a given target by a random basis is amenable to methods of statistical physics such as replica theory [1]. In particular, one can compute the expected typical fitness of a random genome (random binary matrices with density *p*) in a close-to-homogeneous environment 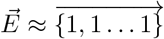 and one finds that the control parameters in the problem are *p*, *α* = *K/L*, and *σ*^2^ = the variance of the components of 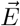. By analogy, for the fitness of a typical genome in two environments 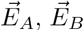 one may expect the control parameters to include the two variances 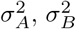 and the component-wise covariance 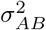. When the genome starts evolving and thus becomes atypical, it is likely that additional features of the environments and/or their relation to each other will become relevant, beyond these basic statistics. In short, whether or how the 2*L*-dimensional space of environment pairs can be reduced to a smaller set of key factors shaping the evolutionary process is not obvious.

Nevertheless, it is clear that the difference between environments is a relevant characteristic. Conveniently, it is also the least model-specific: any model of two-environment evolution and any experimental setup will have a natural counterpart. At the same time, however we define Δ*E*, the argument above demonstrates that it will never be the *only* relevant feature: even assuming 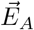 and 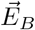 are drawn from the same ensemble (so that *σ*_*A*_ = *σ*_*B*_), we still expect the space of control parameters to be at least two-dimensional (*σ*_*A*_ and *σ_AB_*). Thus, we do not necessarily expect evolution in two environment pairs to look statistically the same simply because they share the same Δ*E*, and the approach of Fig. 3 grouping random environment pairs based solely on their Δ*E* is only an educated guess. Fortunately, for the coarse characterization of the dynamics of *χ* this approach proved sufficient: the standard deviation over 100 replicates (shaded on the plot) is remarkably small.

These considerations provide the additional motivation behind our approach of “exaggerating” or “softening” the differences in a given (randomly chosen) pair. This procedure changes Δ*E* while preserving much of the structure of the two environments in relation to each other that could be relevant for evolution, such as whether the largest or smallest entries of 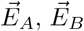 align with each other, etc. As a result, in Fig. 4 and 5, evolving a genome in an environment pair 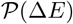 with an exaggerated or softened Δ*E* tends to also increase its fitness in the original pair of interest 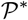.

### S2. QUANTIFYING TRADEOFFS

In this work, we largely focused on a single metric for quantifying tradeoffs, namely the mutational tradeoff *χ*:

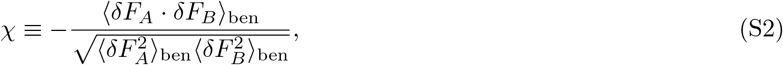

where *δF*_*A*_, *δF*_*B*_ are the two fitness effects of each mutation, and the averaging omits doubly-deleterious mutants. This definition was adopted for its three key strengths:

1. Mutational tradeoff *χ* is a property of a single genome (not a population of genomes), simplifying simulations;
2. Computing *χ* requires only a *local* knowledge of the fitness landscape (the immediate vicinity of the genome considered);
3. Mutational tradeoff is directly related to expectations for near-term evolutionary future of a genome (cf. Lande’s theory of selection on correlated traits [2]).

Let us also highlight some of its caveats. First, our definition is “scale-invariant”: scaling *δF*_*A*_ and *δF*_*B*_ by the same positive factor leaves *χ* intact. In other words, the mutational tradeoff *χ* characterizes only the shape of the mutant cloud, and is not sensitive to the absolute magnitude of the underlying fitness effects. In many respects, this is an advantage. However, it does mean that judging the relevance of *χ* changing along an evolutionary trajectory requires an independent verification that the underlying fitness effects are not negligible. In particular, when interpreting the dynamics of *χ* observed in Fig. 3A we must reassure ourselves that the genomes we are tracking are not running out of beneficial mutations. This reassurance is provided by Fig. 5A, which confirms that genomes continue evolving in fitness throughout the considered time period. In Fig. 4 the reassurance is automatic, as we are plotting *χ* against fitness, rather than time.

The second caveat we must highlight is that mutant clouds do not always look like the examples shown in Fig. 2B. A representative cross-section is shown in Fig. S2. The figure draws on the extensive simulations performed for Fig. 3 in the main text to display randomly selected examples with mutational tradeoff strength scoring close to −1, 0 or 0.5. In each group, examples are ordered left to right by time at which they were observed. Note the two highlighted examples where the value of *χ* is not informative for reasons we just discussed, because the beneficial mutations are too weak / too few (blue “x”). However, overall, at the timescales investigated here (~ 250 mutation steps), the genomes do not exhibit any significant depletion of available mutations (as they would at later stages of evolution; compare to Fig. S1). Note also some, but not all of the middle panels exhibit the “cross shape” characteristic of a modular architecture (cf. Fig. 2B). This point is further discussed below, section “Mutational modularity is not equivalent to *χ* = 0”.

**FIG. S2.**
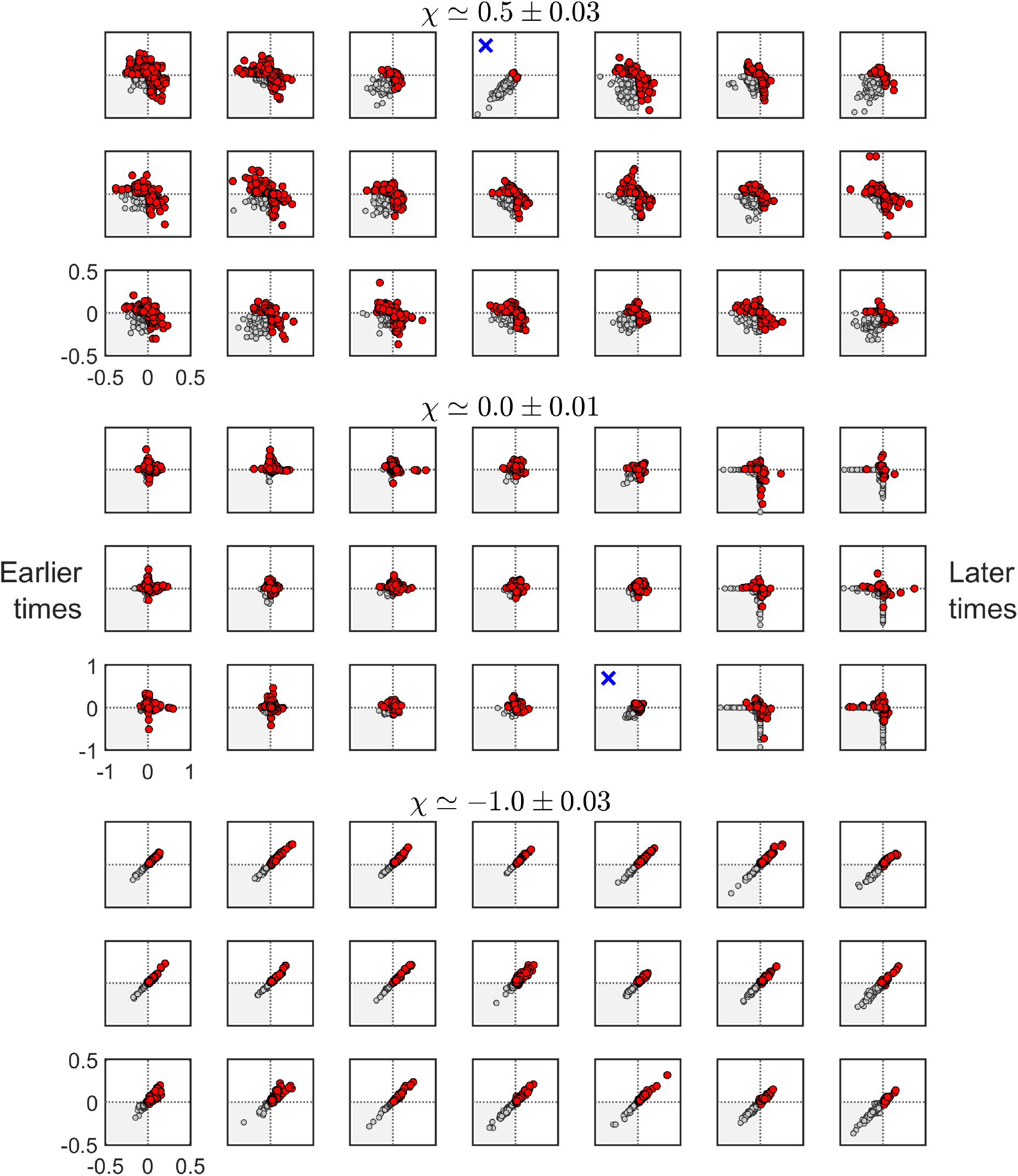
Mutant clouds for examples genomes grouped by their mutational tradeoff score *χ* (compare to Fig. 2B); within each group, examples are ordered by timepoint when observed (earlier timepoints on the left, later timepoints on the right). Individual panels as in Fig. 2B: only mutations beneficial in at least one of the environments (in red) contribute to the tradeoff score; doubly deleterious mutations in gray are ignored. Shown are 21 example per group; in each group axis ranges are the same; labels not shown to reduce clutter. In two cases (identified here by eye, and marked with a blue x) with only a few beneficial mutations available, the mutational tradeoff score is unreliable. At *K* = 4 and *N* = 100 (400 datapoints in each mutant cloud), and for early stages of evolution (Fig. S1), such cases are rare.

#### Other metrics for quantifying tradeoff strength

The metric we use (mutational tradeoff) is certainly not the only way to quantify the notion of a “tradeoff strength”. For example, another perfectly reasonable approach might be to consider the *population-level tradeoff* : sampling extant organisms, and asking whether those better at task *A* tend to be worse at task *B*, and whether perhaps the observed phenotypes trace out a Pareto front one would associate with a tradeoff [3, 4].

Specifically in the context of our model, imagine drawing 100 random starting genomes, and consider the scatter point of their fitness values in the two environments, both initially and as a function of time as these genomes are independently evolved (Fig. S3A). (In other words, rather than sampling organisms in a population, we sample multiple independently evolving populations, but we will still call it “population tradeoff” for conciseness.) We observe that initially, the dominant axis of variation was that some genomes were better at both tasks, while other were worse at both tasks. After 250 mutation steps, the pattern is transformed into an anti-correlation: genomes better in environment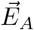 tend to be worse in 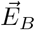. The degree of anticorrelation of this evolving cloud can also be said to capture the intuitive notion of a performance tradeoff, and is distinct from the mutational tradeoff *χ* considered in the main text.

Fig. S3B shows the population tradeoff as a function of time, computed across 100 independently evolving populations from 100 random starting points, evolving at different Δ*E* (but unlike the setting of Fig. 3, now the environment pair is of course the *same* for all 100 replicates). This figure should be compared with Fig. 3A in the main text. We see that the evolution of this measure of tradeoff exhibits the exact same qualitative trends discussed in the main text, namely: (1) for random genomes (timepoint 0), tradeoff is strongest when Δ*E* is largest; (2) tradeoff strength generally increases with time; and (3) for genomes evolving at a large Δ*E*, the initial increase is followed by a decline in tradeoff strength. The figure confirms that the observations of Fig. 3A in the main text are not an artifact of the one definition on which we chose to focus.

The curve Δ*E* = 0.1 in Fig. S3B deserves a special comment. Just like for Δ*E* = 0.5 (panel A), as mutational tradeoff becomes strong, the initial cloud (strongly diagonally elongated) starts to “bunch up” (genomes at the front stall, the ones at the back catch up) and will eventually become antidiagonal. But since at Δ*E* = 0.1 the random initial genomes start out very correlated in fitness, for Δ*E* = 0.1 this bunching-up process takes much longer. By generation ~ 150, individually, each of the evolving genomes is already facing a strong mutational tradeoff (as demonstrated in Fig. 3A in the main text). However, at the population level, some genomes are still ahead of others in both environments. While this tradeoff will eventually propagate to the population level, there is an extended transient during which the population-level fitness remains correlated (mutational tradeoff is masked). This is why, in the main body of the paper, we chose to focus on the mutational tradeoff, as it is the first to change, and is also the property most directly linked to shaping evolutionary future. Conveniently, its evaluation is also much less costly computationally, as it is a property of a single genome and not a population. Still, the nontrivial relation and interplay between the local (mutational) picture and the global (observed across extant organisms) is extremely interesting, and the “masking effect” described above becomes particularly important when trying to infer tradeoffs from experimental observations [5].

**FIG. S3.**
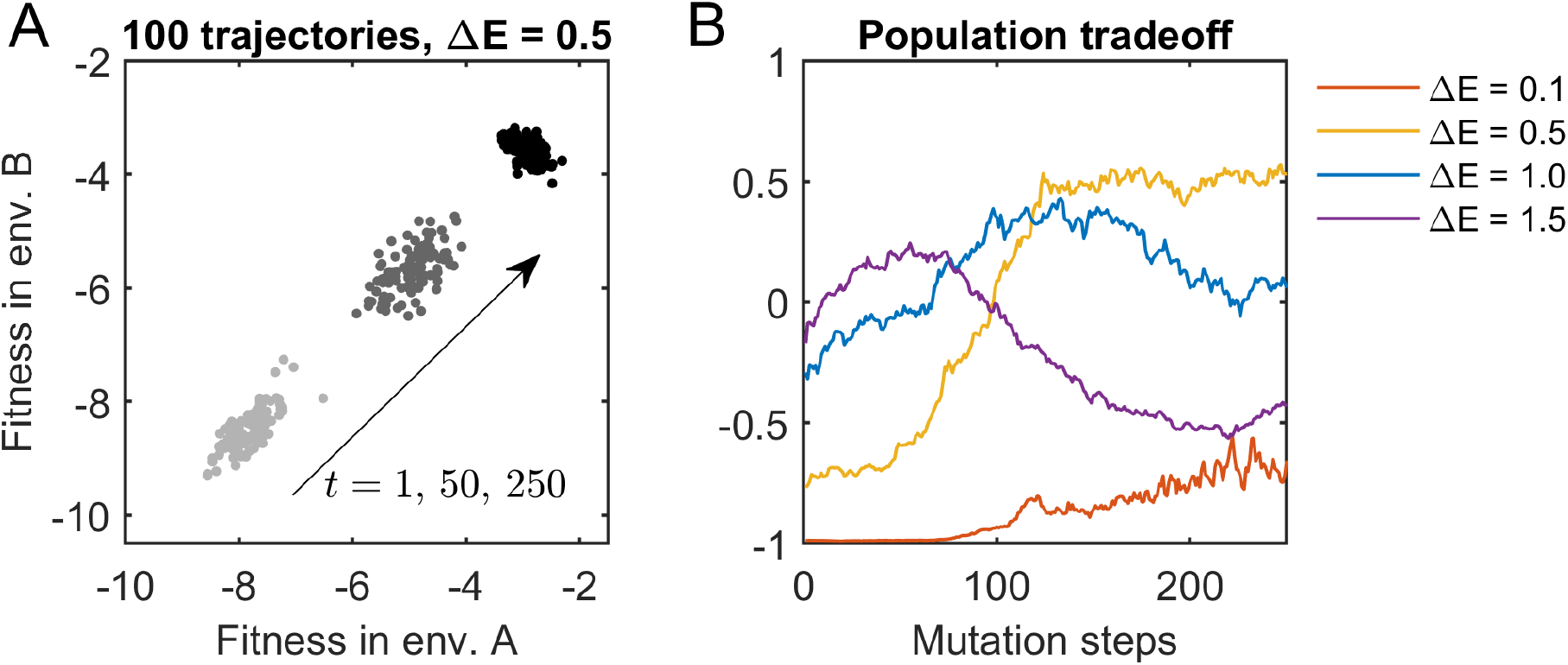
A different definition of tradeoff strength exhibits the same qualitative behavior of initial increase in tradeoff strength and a subsequent decrease if evolving at large Δ*E* (compare to Fig. 3A in the main text). Left: Define population-level tradeoff as the degree of anticorrelation of the fitness values across a population of 100 genomes evolving independently from random starting points. Shown is the scatter plot for the entire simulated population at 3 time points. Right: The analog of Fig. 3A, showing the evolution of population-level tradeoff strength as a function of time for a population evolving at different Δ*E*.

Beyond these considerations, yet another approach to quantifying tradeoffs might be in terms of the “cost of being a generalist”: to what extent does evolving a genome under pressure from both 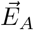 and 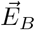 hurt its performance, compared to what would be achievable by evolving in either environment alone? Such questions are also addressable in our framework, but are left for future work.

Ultimately, no strategy of characterizing tradeoff strength by a single number could adequately capture this multifaceted notion. The approach adopted here is meant as a reasonable starting point to illustrate the theoretical framework we present.

### S3. MODULARITY

#### Two definitions of modularity

The term “modularity”, while widely used, lacks a consensus definition. In our context, one could in fact define modularity in *two* distinct ways. First, as a mutational property: a perfectly modular genome is one where a mutation improving performance in one environment does not affect the other. Alternatively, modularity could refer to the usage of regulators across environments: a modular genome is one where each regulator (basis vector) is specialized to (only used in) one or the other environment. We stress that in either case, modularity is defined in reference to a particular set of environments.

The two definitions are related, but distinct. If a given basis vector is perfectly specialized, a mutation in it will only affect performance in one environment but not the other. However, for weakly modular genomes, one might imagine either property arising first. The ability to define modularity in both ways, allowing the relation between them to be characterized, is one of the advantages of a framework like ours, where expression coefficients are both explicit and environment-dependent. However, for the purposes of this work, we use the term “modularity” exclusively in its mutational sense above, because (a) this definition is measurable experimentally, and (b) the mutational perspective on modularity is the one most obviously linked to the notion of “evolvability”, allowing us to comment on this relation (and nuance it) in Fig. 4D, E.

#### Mutational modularity is not equivalent to *χ* = 0

It is important to stress that although mutational modularity in the sense described above necessarily entails *χ* ≈ 0, the converse is not true: as illustrated in Fig. 2B, a genome with *χ* ≈ 0 is not necessarily modular. We can see this in the middle panels of Fig. S2, some, but not all of which exhibit the cross shape characteristic of mutational modularity (Fig. 2B).

For this reason, the weakening of mutational tradeoff cannot be directly interpreted as the emergence of modularity, unless additional analysis is performed, as we did in Fig. 3. To avoid confusion, in the main text we preferentially use the former phrasing.

